# Xenium-based spatial transcriptomic screening identifies candidate mRNAs localized in neuronal and glial processes of the adult mouse cerebellum

**DOI:** 10.64898/2026.06.11.731766

**Authors:** Sotaro Ito, Toma Adachi, Kyoka Suyama, Masaki Sone, Mikio Hoshino

## Abstract

Subcellular mRNA localization contributes to local protein synthesis and functional compartmentalization in polarized cells, including neurons and glial cells. Although process-localized mRNAs have been identified by *in situ* hybridization, reporter-based analyses, and compartment-based transcriptomic approaches, systematic screening of such mRNAs within intact brain tissue while preserving tissue architecture remains challenging. Here, we developed a Xenium-based spatial transcriptomic screening strategy to identify candidate mRNAs localized in neuronal and glial processes in the adult mouse cerebellum. We focused on the molecular layer, which is densely occupied by Purkinje cell dendrites, Bergmann glial radial processes, and granule cell parallel fibers, but contains relatively few cell bodies. Using a Xenium Prime 5K dataset from adult mouse cerebellar sections, we first identified 199 genes whose transcripts were enriched in the molecular layer. By further focusing on DAPI-negative, process-rich regions and reducing contributions from molecular layer cell bodies, we extracted 126 candidate process-localized genes. Comparison with Allen Brain Atlas *in situ* hybridization data supported molecular layer localization with process-like patterns for 28 of 32 evaluable candidates. Integration with a published adult cerebellar single-nucleus RNA-seq atlas provided information on the possible cellular origins of these candidates. Gene Ontology analysis revealed enrichment of terms related to intracellular transport, cell projection structures, and synaptic function. These findings support the usefulness of a Xenium-based spatial transcriptomic first-screening strategy for identifying candidate process-localized mRNAs in intact brain tissue and provide a resource for studying RNA localization in cerebellar neuronal and glial processes.

## Introduction

Subcellular mRNA localization is an important mechanism by which polarized cells spatially regulate protein synthesis and cellular function (Buxbaum et al., 2015; Holt and Schuman, 2013; Meservey et al., 2021). Although many mRNAs are enriched in the soma, a subset of transcripts is transported to distal subcellular compartments, including dendrites, axons, growth cones, synaptic compartments, and glial processes (Doyle and Kiebler, 2011; Holt and Schuman, 2013; Buxbaum et al., 2015; Cioni et al., 2018; Mofatteh, 2020; Turner-Bridger et al., 2020; Dalla Costa et al., 2021). In neurons, local translation of these mRNAs contributes to axon guidance, synapse formation, synaptic plasticity, neuronal maintenance, and circuit function (Lin and Holt, 2007; Jung et al., 2011; Holt and Schuman, 2013; Ohashi and Shiina, 2020). Although local mRNA translation has been studied most extensively in neurons, glial cells also possess complex and functionally specialized processes, and increasing evidence suggests that mRNA transport and local translation contribute to glial development, morphology, intercellular interactions, and responses to neuronal activity (Sakers et al., 2017; Barton et al., 2019; Meservey et al., 2021).

Previous studies have identified process-localized mRNAs using several approaches. In situ hybridization (ISH), reporter-based analyses, and single-gene imaging approaches have been useful for examining the subcellular localization of specific transcripts (Doyle and Kiebler, 2011; Xing and Bassell, 2013; Lee et al., 2016). In addition, compartmentalized culture systems, microfluidic chambers, and biochemical isolation of neurites, dendrites, or axons have enabled transcriptome-wide identification of mRNAs enriched in neuronal processes by RNA sequencing (Taylor et al., 2009; Gumy et al., 2011; Zappulo et al., 2017; Middleton et al., 2018; Dargan et al., 2025). These approaches provide either high-resolution information for selected transcripts or broad transcriptomic information from physically isolated subcellular compartments. However, they often require dissociated or cultured cells, reporter systems, or mechanical separation of cellular compartments, which can disrupt native tissue architecture, cell-cell interactions, and the anatomical context in which cellular processes normally exist. Thus, methods for systematically screening candidate process-localized mRNAs within intact brain tissue remain insufficiently established.

Recent advances in spatial transcriptomics provide new opportunities to address this issue. High-plex in situ platforms such as Xenium In Situ can detect individual RNA molecules corresponding to many target genes directly on tissue sections with subcellular spatial resolution while preserving tissue architecture (Williams et al., 2022; Janesick et al., 2023). Because individual RNA molecules are detected as spatially resolved spots, these technologies make it possible to examine not only which cell types express particular transcripts, but also where those transcripts are positioned within tissue and subcellular contexts. This feature is particularly useful for analyzing brain regions in which cell bodies and cellular processes are spatially segregated. Therefore, by selecting an appropriate anatomical region, Xenium-based spatial transcriptomic analysis may be useful as a first-screening strategy for identifying candidate process-localized mRNAs within intact nervous tissue.

The mature mammalian cerebellar cortex provides a suitable anatomical system for this purpose. It is organized into three major layers: the molecular layer (ML), Purkinje cell layer (PCL), and inner granule cell layer (IGL) (Leto et al., 2016). The ML contains relatively few cell bodies, including molecular layer inhibitory neurons (ML-INs) and rare glial lineage cells (Consalez et al., 2013; Sotelo, 2015; Singer et al., 2025), but is densely occupied by cellular processes. These include Purkinje cell (PC) dendrites, Bergmann glial (BG) radial processes, and granule cell (GC) axons known as parallel fibers (Ango et al., 2008; De Zeeuw and Hoogland, 2015; Leto et al., 2016). BGs are highly polarized astroglial cells whose somata are located in or near the PCL and whose radial processes extend through the ML toward the pial surface, where they interact closely with PC dendrites and synaptic structures. Therefore, the cerebellar ML offers a unique tissue environment in which mRNA signals detected outside DAPI-positive cell body regions are likely to include transcripts localized in neuronal and glial processes.

In this study, we developed a Xenium-based spatial transcriptomic screening strategy to identify candidate process-localized mRNAs in the adult mouse cerebellar ML. We reanalyzed a Xenium Prime 5K dataset generated from FFPE cerebellar sections of an 8-month-old mouse and extracted genes whose transcripts were enriched in DAPI-negative, process-rich regions of the ML. This screening identified 126 candidate process-localized genes. We then evaluated these candidates by comparison with Allen Brain Atlas ISH data, inferred their possible cellular origins using a published adult mouse cerebellar single-nucleus RNA-seq atlas, and examined their functional characteristics by Gene Ontology (GO) analysis. Our findings suggest that Xenium-based spatial transcriptomics can serve as a useful first-screening approach for identifying candidate process-localized mRNAs while preserving intact brain tissue architecture.

## Results

### Xenium-based extraction of candidate process-localized mRNAs in the adult mouse cerebellar ML

To identify candidate process-localized mRNAs within intact brain tissue, we focused on the molecular layer (ML) of the adult mouse cerebellum, a process-rich region containing relatively few cell bodies. The adult cerebellar ML is densely occupied by Purkinje cell (PC) dendrites, Bergmann glial (BG) radial processes, and granule cell (GC) parallel fibers, whereas only a small number of cell bodies, including molecular layer inhibitory neurons (ML-INs), are present. We reasoned that this anatomical organization would provide a useful tissue context for spatially screening mRNA signals associated with neuronal and glial processes.

For this analysis, we reanalyzed a Xenium Prime 5K spatial transcriptomic dataset that was previously generated by our group from FFPE cerebellar sections of an 8-month-old ICR mouse and deposited in Zenodo as part of Suyama et al. (in press) (Figure 1a; DOI: 10.5281/zenodo.16562843). In this dataset, mRNA signals corresponding to 5,006 target genes included in the Xenium Prime 5K Mouse Pan Tissue and Pathways Panel were detected as spatially resolved spots on tissue sections. We first examined the spatial distribution of DAPI signals, *18S* rRNA signals, and individual mRNA spots. The laminar structure of the cerebellar cortex, including the ML, Purkinje cell layer (PCL), and inner granule cell layer (IGL), was clearly distinguishable. Numerous mRNA spots were detected not only in the PCL and IGL, but also in the ML (Figure 1a, Figure S1a, b).

**Figure 1.**
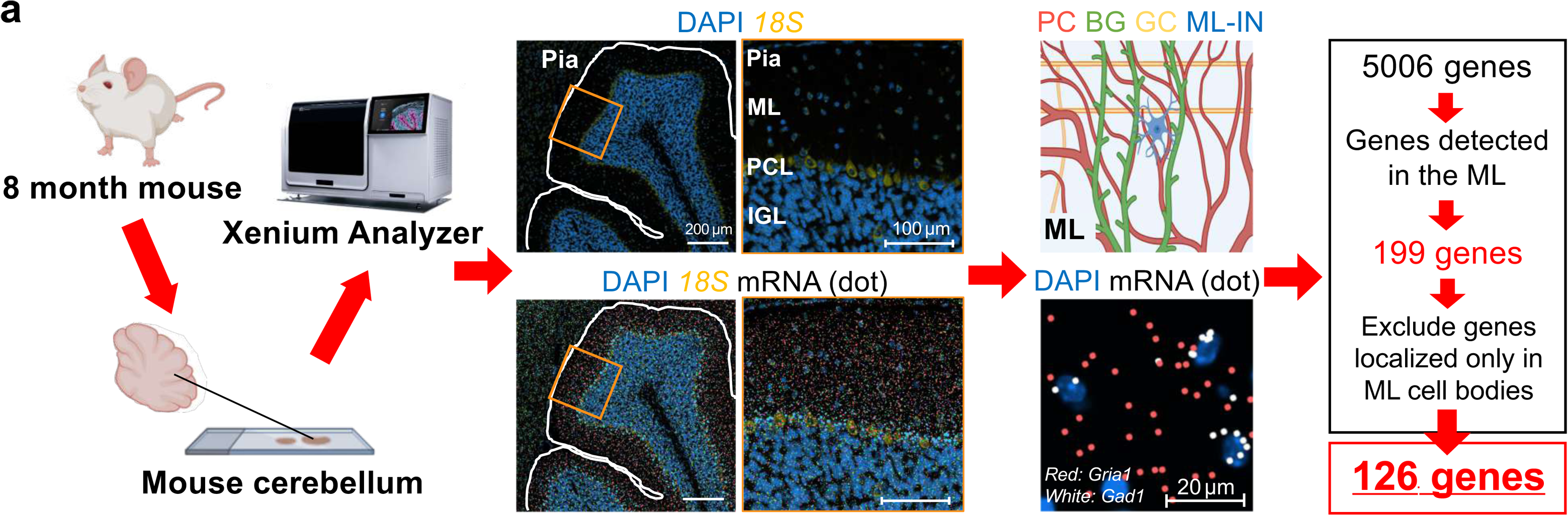
Xenium-based screening of candidate process-localized mRNAs in the adult mouse cerebellar molecular layer. (a) Schematic overview of Xenium Prime 5K spatial transcriptomic analysis using cerebellar sections from an 8-month-old ICR mouse and the workflow for extracting candidate process-localized mRNAs detected in the adult cerebellar molecular layer (ML). The laminar structure of the cerebellar cortex was identified using DAPI and *18S* rRNA signals, and the spatial distribution of individual mRNA spots was analyzed. The adult cerebellar ML is densely occupied by Purkinje cell (PC) dendrites, Bergmann glial (BG) radial processes, and granule cell (GC) parallel fibers, whereas only a small number of cell bodies, including those of molecular layer inhibitory neurons (ML-INs), are present. First, genes detected in the ML at a density of 1,500 dots/mm² or higher were extracted, yielding 199 genes. Next, to exclude genes localized only in ML cell bodies, including ML-INs, genes detected in DAPI-negative regions of the ML at a density of 1,200 dots/mm² or higher were extracted, resulting in the selection of 126 genes as candidate process-localized genes. In the magnified image at the lower right, *Gria1* mRNA spots are shown in red and *Gad1* mRNA spots are shown in white. Scale bars: 200 µm, 100 µm, and 20 µm.

As a first-pass screen for ML-detected transcripts, we selected genes whose transcripts were detected in the ML at a density of 1,500 dots/mm² or higher. This criterion identified 199 genes (Figure 1a). However, ML-wide enrichment alone cannot distinguish process-associated transcripts from transcripts present in sparse ML cell bodies. Indeed, the ML contains a small number of cell bodies, including *Gad1*-positive ML-INs and rare glial lineage cells (Consalez et al., 2013; Sotelo, 2015; Singer et al., 2025), and some mRNA spots detected in the ML appeared to be associated with DAPI-positive regions or their immediate vicinity (Figure S1c).

To enrich for transcripts associated with neuronal and glial processes, we next focused on DAPI-negative regions within the ML. These regions are expected to be enriched in PC dendrites, BG radial processes, and GC parallel fibers, while being depleted of somatic RNA signals. Genes detected in these DAPI-negative ML regions at a density of 1,200 dots/mm² or higher were selected as candidate process-localized genes. This second screening step identified 126 genes (Figure 1a, Figure 2, Figure S2). Hereafter, we refer to these 126 genes as candidate process-localized genes and to their transcripts as candidate process-localized mRNAs. Thus, this two-step workflow narrowed the 5,006 genes included in the Xenium panel to 126 candidate process-localized genes.

**Figure 2.**
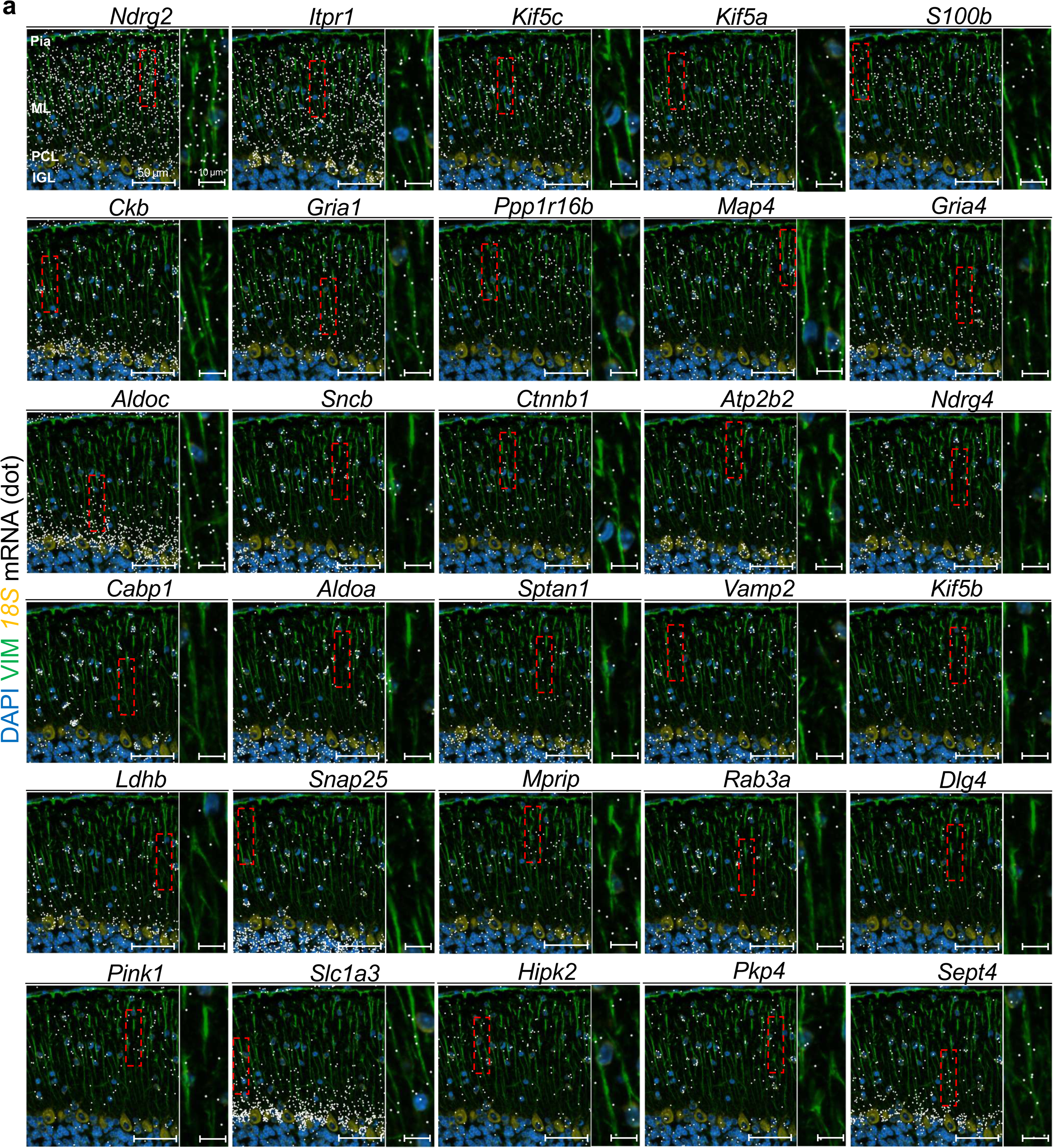
Representative Xenium images of candidate process-localized mRNAs in the adult cerebellar molecular layer. (a) Representative Xenium images showing candidate process-localized mRNAs detected in the adult mouse cerebellar molecular layer (ML). mRNA spots for each indicated gene are shown in white. DAPI, VIMENTIN (VIM), and *18S* rRNA signals are shown in blue, green, and orange, respectively. Red dashed boxes indicate representative ML regions where mRNA spots were detected among densely distributed VIM-positive processes. Magnified images of the boxed regions are shown to the right of each panel. Additional candidate genes are shown in Figure S2. Scale bars: 50 µm and 10 µm.

To visually assess the spatial distribution of these candidates, we next examined representative Xenium images. mRNA spots for multiple candidate process-localized genes, including *Ndrg2*, *Itpr1*, *Kif5c*, *Kif5a*, *S100b*, *Ckb*, *Gria1*, *Slc1a3,* and others, were broadly detected in the ML, including DAPI-negative regions (Figure 2, Figure S2). Consistent with the dense distribution of BG radial processes in the ML, some candidate mRNA spots appeared to overlap with VIMENTIN (VIM)-positive BG process-like structures. These observations support the idea that the extracted candidates include mRNAs present in process-rich regions of the ML. Together, these results suggest that Xenium analysis of DAPI-negative, process-rich regions of the adult cerebellar ML can serve as a first screening strategy to extract candidate process-localized mRNAs while preserving tissue architecture.

### Evaluation of candidate process-localized mRNAs using Allen Brain Atlas ISH data

To obtain independent support for the Xenium-based extraction of candidate process-localized mRNAs, we compared the spatial distribution of transcripts encoded by the 126 candidate process-localized genes with publicly available *in situ* hybridization (ISH) data from the Allen Brain Atlas. The Allen Brain Atlas provides ISH images for many genes in the mouse brain and is useful for assessing whether candidate transcripts show consistent anatomical localization patterns in the cerebellum (Lein et al., 2007; Sunkin et al., 2013). Because P28 cerebellar ISH images were available for the largest number of candidate genes, we used this dataset to evaluate broad ML localization patterns.

Among the 126 candidate process-localized genes, cerebellar ISH images suitable for evaluation were available for 32 genes. We classified genes as supported when the available ISH images showed clear signals in the ML with process-like staining patterns. Among the 32 evaluable candidates, 28 genes showed clear ML signals with process-like patterns in the Allen Brain Atlas ISH images (Figure 3). In contrast, the remaining four genes did not show clear ML localization or process-like patterns in the available ISH images (Figure S3).

**Figure 3.**
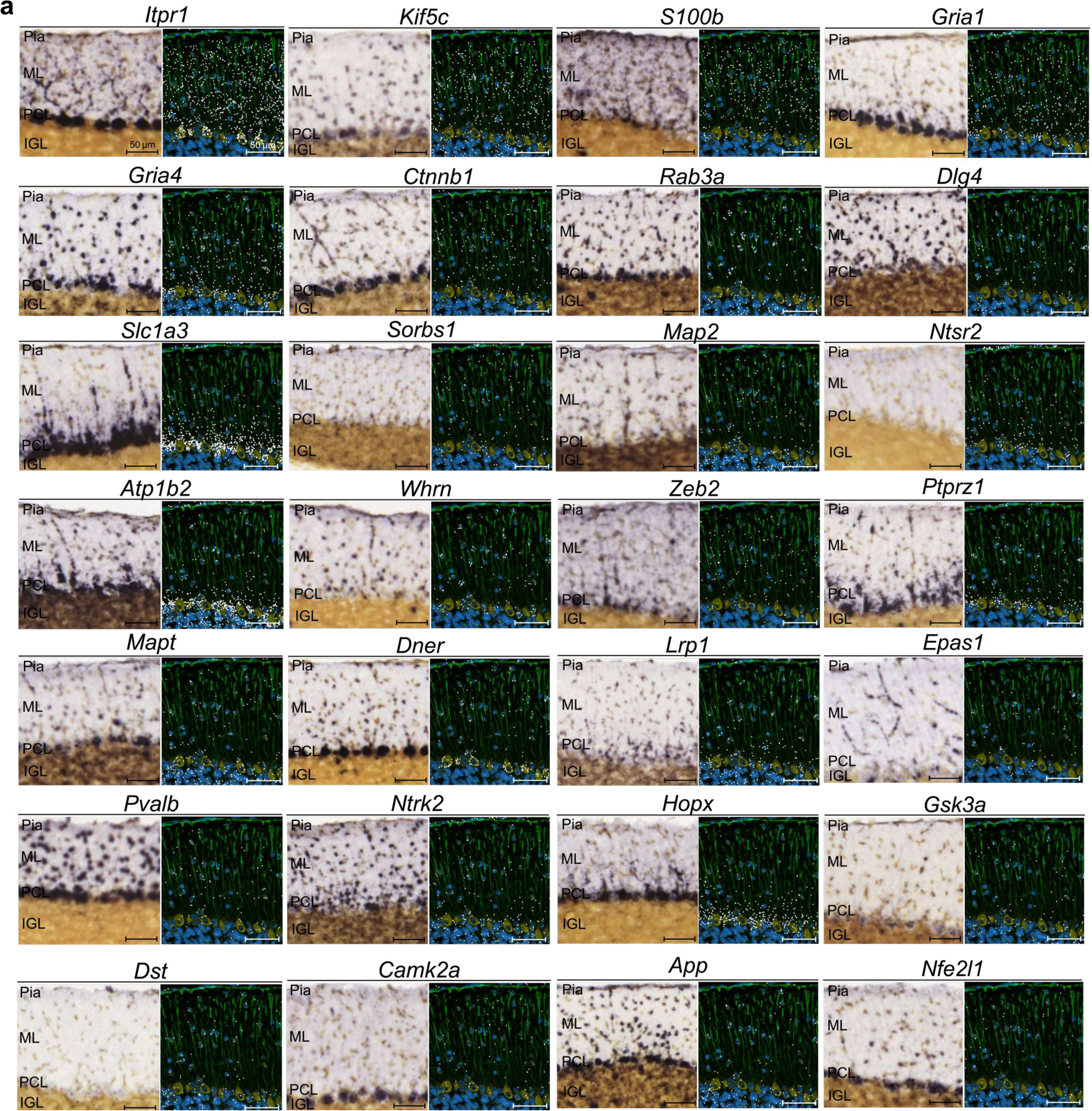
Comparison with Allen Brain Atlas ISH data supports ML localization of candidate process-localized mRNAs. (a) Representative comparison of Allen Brain Atlas *in situ* hybridization (ISH) images and Xenium images for 28 candidate process-localized genes that showed clear ISH signals in the molecular layer (ML) with process-like staining patterns. For each gene, the Allen Brain Atlas ISH image is shown on the left, and the corresponding Xenium image is shown on the right. In the Xenium images, mRNA spots for each indicated gene are shown in white. DAPI, VIMENTIN (VIM), and *18S* rRNA signals are shown in blue, green, and orange, respectively. Scale bars: 50 µm.

Thus, 87.5% of the evaluable candidates (28/32) were supported by independent ISH data. These observations support the ability of our Xenium-based screening strategy to enrich for genes whose transcripts show ML-associated, process-like distribution patterns. Together with the Xenium images shown above, the Allen Brain Atlas comparison provides independent anatomical support for the 126 candidate process-localized mRNAs extracted from DAPI-negative, process-rich regions of the adult cerebellar ML.

### Inference of the cellular origin of candidate process-localized mRNAs using a published snRNA-seq atlas

Next, to infer the possible cellular origin of the 126 candidate process-localized mRNAs, we examined the cell-type-specific expression patterns of their corresponding genes using a published adult mouse cerebellar snRNA-seq atlas (Kozareva et al., 2021) (Figure 4, Figure S4). Because the adult cerebellar ML is densely occupied by PC dendrites, BG radial processes, and GC parallel fibers (Ango et al., 2008; De Zeeuw and Hoogland, 2015; Leto et al., 2016), we focused on PC, BG, and GC clusters in this atlas. For this analysis, we operationally defined genes with a median unique molecular identifier (UMI) count of 1.0 or higher in a given cell-type cluster as being expressed in that cell type.

**Figure 4.**
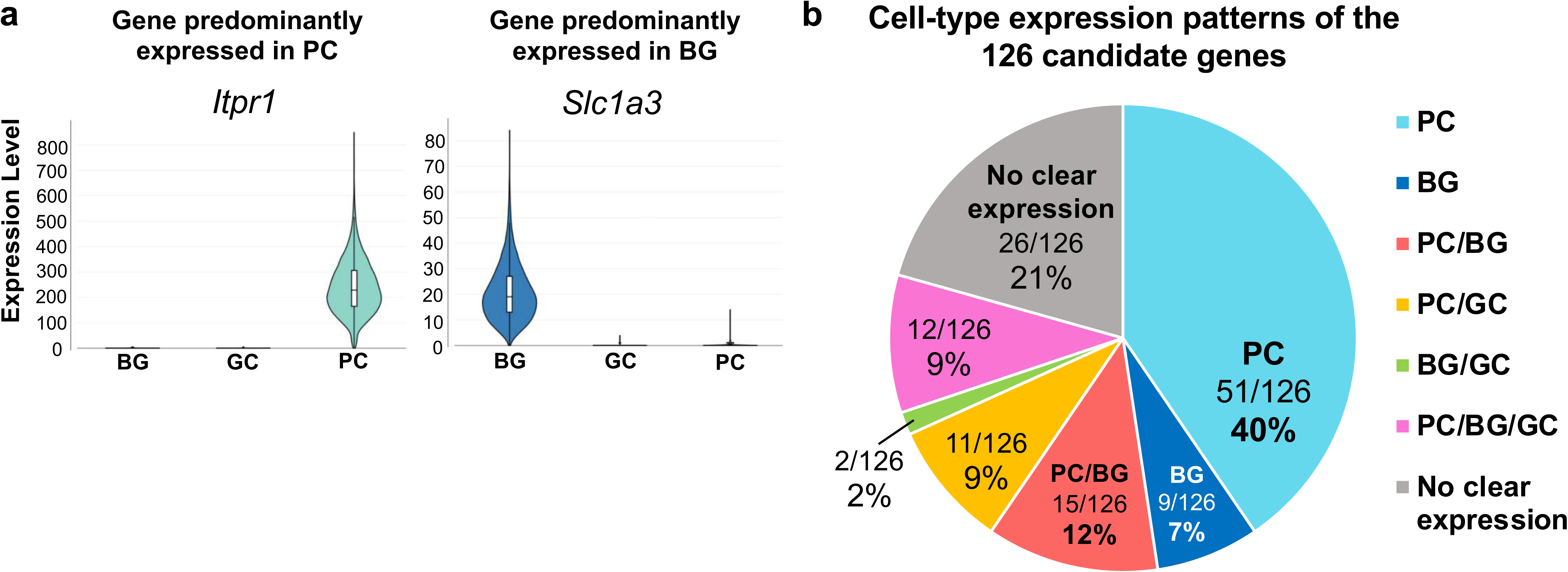
Cell-type expression analysis of candidate process-localized genes using a published cerebellar snRNA-seq atlas. (a) Representative violin plots showing cell-type-specific expression of candidate process-localized genes in the adult mouse cerebellar single-nucleus RNA-seq (snRNA-seq) atlas reported by Kozareva et al. *Itpr1* is shown as a representative gene predominantly expressed in Purkinje cells (PCs), and *Slc1a3* is shown as a representative gene predominantly expressed in Bergmann glial cells (BG). Expression levels are shown as unique molecular identifier (UMI) counts in BG, granule cell (GC), and PC clusters. (b) Summary of cell-type expression patterns of the 126 candidate process-localized genes based on the published cerebellar snRNA-seq atlas. Genes with a median UMI count of 1.0 or higher in a given cell-type cluster were considered to be expressed in that cell type. Candidate genes were classified into the following categories: PC only, BG only, PC/BG, PC/GC, BG/GC, PC/BG/GC, and no clear expression in BG, GC, or PC clusters. The number and percentage of genes in each category are indicated.

As representative examples, *Itpr1* showed strong expression in PC clusters, whereas *Slc1a3* showed preferential expression in BG clusters (Figure 4a). We then classified the 126 candidate process-localized genes according to their expression patterns in PC, BG, and GC clusters. The largest fraction consisted of genes expressed only in PCs, with 51 genes (40.5%) falling into this category (Figure 4b, Figure S4a, b). Genes expressed only in BGs accounted for 9 genes (7.1%; Figure 4b, Figure S4c). No genes met the expression threshold only in GCs (Figure 4b). In addition, 15 genes were expressed in both PCs and BGs (11.9%; Figure 4b, Figure S4d), 11 genes in both PCs and GCs (8.7%; Figure 4b, Figure S4e), 2 genes in both BGs and GCs (1.6%; Figure 4b, Figure S4f), and 12 genes in all three cell types (9.5%; Figure 4b, Figure S4g). The remaining 26 genes did not meet the expression threshold in any of the PC, BG, or GC clusters (20.6%; Figure 4b, Figure S4h).

These results suggest that many candidate process-localized mRNAs extracted from the ML are likely derived from PCs, consistent with the dense arborization of PC dendrites in the ML. Importantly, a subset of candidates was expressed in BGs alone or in BGs together with PCs and/or GCs, suggesting that the screening also captured candidates potentially associated with Bergmann glial processes. Thus, integration of Xenium-based spatial screening with a published snRNA-seq atlas enabled inference of the possible cellular origins of candidate process-localized mRNAs.

### Functional characterization of candidate process-localized genes by GO analysis

Finally, to characterize the functional features of the 126 candidate process-localized genes, we performed Gene Ontology (GO) enrichment analysis using ShinyGO (Ge et al., 2020), with the 5,006 genes included in the Xenium panel used as the background gene set (Figure 5).

**Figure 5.**
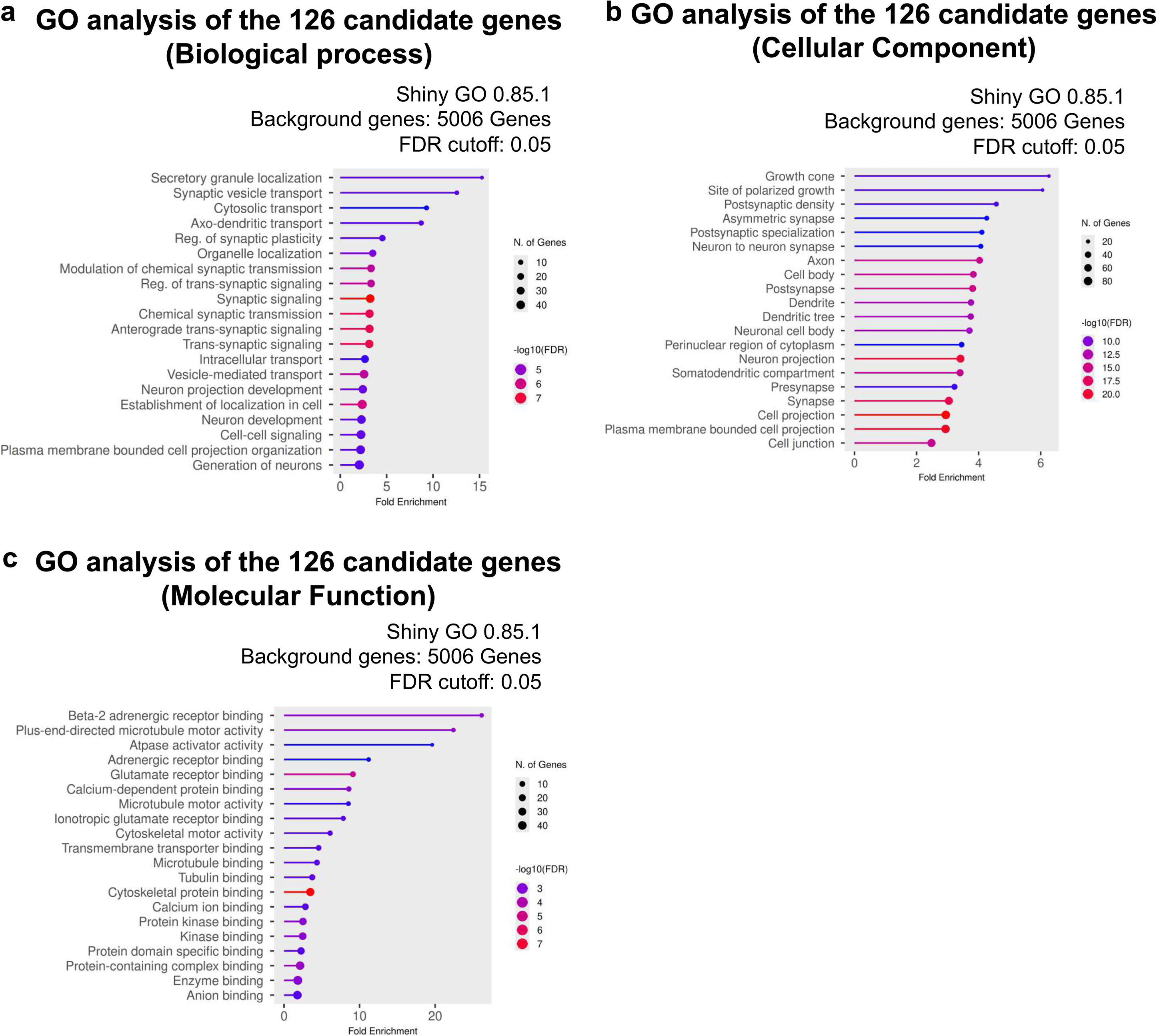
Functional characterization of candidate process-localized genes by GO analysis. (a–c) Gene Ontology (GO) enrichment analysis of the 126 candidate process-localized genes extracted from Xenium data. GO analysis was performed using ShinyGO v0.85.1 with 5,006 genes included in the Xenium Prime 5K dataset as background genes and an FDR cutoff of 0.05. Enriched GO terms are shown for Biological Process (a), Cellular Component (b), and Molecular Function (c). Dot size indicates the number of genes assigned to each GO term, and color indicates −log10(FDR). The x-axis indicates fold enrichment.

In the Biological Process category (Figure 5a), enriched terms included those related to intracellular and process-associated transport, such as secretory granule localization, synaptic vesicle transport, cytosolic transport, and axon-dendritic transport. Terms related to synaptic regulation, including regulation of synaptic plasticity, were also enriched. In the Cellular Component category (Figure 5b), enriched terms were associated with cellular projections and synaptic compartments, including growth cone, site of polarized growth, postsynaptic density, asymmetric synapse, postsynaptic specialization, axon, dendrite, and cell projection. In the Molecular Function category (Figure 5c), enriched terms included microtubule motor activity, cytoskeletal motor activity, glutamate receptor binding, transmembrane transporter binding, and related functions.

These GO enrichment patterns indicate that the 126 candidate genes are enriched for functional categories related to intracellular transport, polarized cellular projections, and synaptic organization. These functional categories are compatible with the possibility that the candidate gene set includes transcripts associated with polarized neuronal and glial processes. Together with the spatial distribution of these candidates in the ML, the Allen Brain Atlas ISH comparison, and the snRNA-seq-based cellular origin analysis, these results further support the utility of this Xenium-based workflow as a first screening strategy for identifying candidate process-localized mRNAs in intact brain tissue.

## Discussion

In this study, we developed a Xenium-based spatial transcriptomic screening strategy to identify candidate process-localized mRNAs in the adult mouse cerebellar molecular layer (ML). The adult mouse cerebellar ML is densely occupied by Purkinje cell (PC) dendrites, Bergmann glial (BG) radial processes, and granule cell (GC) parallel fibers, but contains relatively few cell bodies. Taking advantage of this anatomical feature, we extracted mRNAs detected in DAPI-negative, process-rich regions of the ML and identified 126 candidate process-localized genes. We further evaluated these candidates by comparing them with existing ISH data registered in the Allen Brain Atlas, inferring their possible cellular origins using a published adult cerebellar single-nucleus RNA-seq (snRNA-seq) atlas, and performing Gene Ontology (GO) enrichment analysis. These results suggest that Xenium-based spatial transcriptomic analysis can be used as a first screening strategy for systematically extracting candidate process-localized mRNAs while preserving native tissue architecture.

A major advantage of this approach is that positional information of individual mRNA spots can be analyzed together with information on anatomical layers and subcellular regions. Conventional ISH and reporter-based analyses are useful for detailed validation of the localization of individual mRNAs (Doyle and Kiebler, 2011; Xing and Bassell, 2013; Lee et al., 2016), but they are not necessarily suitable for candidate screening across many genes. On the other hand, RNA-seq analysis of isolated neuronal processes in cultured neurons can identify mRNAs enriched in processes (Middleton et al., 2018; Dargan et al., 2025), but this approach loses information on tissue architecture and cellular arrangement. In contrast, Xenium data allow simultaneous evaluation of DAPI, *18S* rRNA, VIM, and the positional information of individual mRNA spots on tissue sections. Therefore, by selecting an anatomical region with relatively few cell bodies and abundant cellular processes, it may be possible to extract candidate process-localized mRNAs while preserving tissue architecture.

The adult cerebellar ML is particularly suitable for this type of analysis because its major cellular components are spatially segregated. The soma of PCs and BGs are located mainly in or near the Purkinje cell layer (PCL), whereas their dendrites or radial processes extend into the ML. In addition, GC soma are located in the inner granule cell layer (IGL), whereas their axons, the parallel fibers, occupy the ML. Therefore, mRNA spots detected in DAPI-negative regions of the ML are likely to include transcripts localized in neuronal and glial processes rather than transcripts confined to soma. This anatomical feature allowed us to use the ML as a tissue-based screening window for candidate process-localized mRNAs. Importantly, the inclusion of BG radial processes in this region makes this system useful not only for studying neuronal mRNA localization, but also for exploring RNA localization in highly polarized glial processes. Although studies of local mRNA regulation have historically focused mainly on neuronal dendrites and axons, glial cells also possess specialized processes that interact closely with neuronal structures. Our approach therefore provides a framework for investigating candidate mRNAs potentially associated with glial processes, including BG radial processes, within their native tissue environment.

Another important aspect of this study is that the candidate genes extracted from Xenium data could be evaluated by integrating multiple public resources. Comparison with ISH data from the Allen Brain Atlas (Lein et al., 2007; Sunkin et al., 2013) provided independent support for ML localization of many evaluable candidates. Although ISH data were available for only a subset of the 126 candidate genes, 28 of 32 evaluable candidates showed clear ML signals with process-like patterns, supporting the idea that the Xenium-based workflow enriched for genes with ML-associated mRNA distributions. In addition, integration with a published adult mouse cerebellar snRNA-seq atlas enabled inference of the possible cellular origins of the candidate genes (Kozareva et al., 2021). This analysis suggested that many candidates are likely transcribed in PCs, consistent with the dense arborization of PC dendrites in the ML, whereas a subset of candidates was expressed in BGs alone or in BGs together with other major cerebellar cell types. Thus, combining spatial transcriptomic screening with cell-type-specific transcriptomic information increases the interpretability of candidate process-localized mRNAs extracted from anatomically defined tissue regions.

GO enrichment analysis using ShinyGO (Ge et al., 2020) provided additional support for the relevance of the candidate gene set to cellular processes and local regulation. The 126 candidate genes were enriched for terms related to intracellular transport, cell projection structures, cytoskeleton-dependent transport, synaptic vesicle transport, postsynaptic specialization, and synaptic function. These categories are consistent with the idea that the extracted gene set includes transcripts associated with polarized cellular compartments such as dendrites, axons, and glial processes. However, GO enrichment should not be interpreted as direct evidence of subcellular localization. Rather, these functional annotations provide an additional layer of evidence suggesting that the Xenium-based screening strategy enriched for genes with biological features compatible with process localization and local regulation.

This study also has several limitations. First, Xenium Prime 5K is a targeted panel and does not cover the whole transcriptome. Therefore, the 126 candidate process-localized genes extracted in this study were selected from genes included in this panel. Second, detection of mRNA spots in DAPI-negative regions does not directly demonstrate that these mRNAs are localized within neuronal or glial processes. Third, evaluation using the Allen Brain Atlas was limited to genes for which public ISH images were available. Fourth, inference of cellular origin using the snRNA-seq atlas was indirect, because snRNA-seq indicates which cell types transcribe each candidate gene but does not directly show which cellular process contains the corresponding mRNA. Therefore, this strategy should be regarded not as a method to conclusively demonstrate process-localized mRNAs, but as a first screening strategy to systematically extract candidates for subsequent validation.

In future studies, candidate genes should be validated individually using high-resolution ISH methods such as RNAscope or smFISH, colocalization analysis with cell-type-specific markers, or functional analysis by genetic manipulation. Nevertheless, the candidate genes extracted in this study included many genes whose ML localization was supported by existing ISH data and genes associated with intracellular transport, synaptic function, and cell projection structures. Thus, Xenium-based spatial transcriptomic analysis may provide a useful first screening strategy for systematically identifying candidate process-localized mRNAs while preserving tissue architecture.

## Experimental procedures

### Xenium dataset

The Xenium dataset analyzed in this study was generated from FFPE cerebellar sections of an 8-month-old ICR mouse and has been deposited in Zenodo (Suyama et al., 2026; DOI: 10.5281/zenodo.16562843). Briefly, 5-µm-thick FFPE brain sections were mounted on Xenium slides and processed using the Xenium Prime 5K Mouse Pan Tissue and Pathways Panel, which included 5,006 genes in the analyzed dataset, according to the manufacturer’s protocol. Deparaffinization, decrosslinking, probe hybridization, ligation, signal detection, cell segmentation, and transcript counting were performed using the Xenium platform and Xenium Analyzer. In the present study, we reanalyzed this adult cerebellar dataset to screen for candidate process-localized mRNAs within the molecular layer (ML).

### Extraction of candidate process-localized mRNAs from the cerebellar molecular layer

The ML, Purkinje cell layer (PCL), and inner granule cell layer (IGL) were identified based on DAPI, *18S* rRNA, and VIMENTIN (VIM) signals, together with the anatomical organization of the cerebellar cortex. Image visualization, region-of-interest (ROI) drawing, and mRNA spot counting were performed using Xenium Explorer v3.2.0.

To identify genes whose transcripts were detected in the ML, ROIs corresponding to the ML were manually drawn in Xenium Explorer, and the number of mRNA spots for each gene within the ML ROI was counted. The density of mRNA spots was calculated as dots/mm². Genes detected in the ML at a density of 1,500 dots/mm² or higher were first extracted as ML-detected genes.

Because the ML contains a small number of cell bodies, including molecular layer inhibitory neurons (ML-INs) (Consalez et al., 2013; Sotelo, 2015; Singer et al., 2025), we further focused on DAPI-negative regions within the ML to enrich for mRNAs detected in process-rich regions. Two square regions of 133 µm × 192 µm were selected within the ML. Within these regions, mRNA spots that did not overlap with DAPI-positive regions were manually counted. Genes detected in DAPI-negative ML regions at a density of 1,200 dots/mm² or higher were selected as candidate process-localized genes, and their transcripts were referred to as candidate process-localized mRNAs.

### Comparison with Allen Brain Atlas ISH data

To evaluate whether the candidate process-localized genes extracted from Xenium data showed independent evidence of ML localization, we compared them with publicly available *in situ* hybridization (ISH) images from the Allen Brain Atlas (Lein et al., 2007; Sunkin et al., 2013). Among the 126 candidate genes, genes for which P28 mouse cerebellar ISH images were available were examined. The P28 dataset was used because it provided the largest number of evaluable ISH images among the available Allen Brain Atlas datasets for the candidate genes. The ML localization pattern of each gene was manually evaluated based on the presence or absence of clear ISH signals in the ML and process-like staining patterns. Genes showing clear ML signals with process-like patterns were considered to be supported by Allen Brain Atlas ISH data (Figure 3), whereas genes without clear ML localization or process-like patterns in the available ISH images were classified separately (Figure S3).

### Cell-type expression analysis using a published cerebellar snRNA-seq atlas

To infer the possible cellular origin of the candidate process-localized mRNAs, we examined their cell-type expression patterns using the published adult mouse cerebellar single-nucleus RNA-seq (snRNA-seq) atlas reported by Kozareva et al. (2021). BG, GC, and PC clusters were selected using the web-based data browser associated with the published atlas (Single Cell Portal; https://singlecell.broadinstitute.org/single_cell/study/SCP795/), and the expression of each candidate gene was queried in these clusters. Genes with a median unique molecular identifier (UMI) count of 1.0 or higher in a given cell-type cluster were considered to be expressed in that cell type. Candidate genes were then categorized according to their expression patterns in BG, GC, and PC clusters.

### GO enrichment analysis

Gene Ontology (GO) enrichment analysis of the 126 candidate process-localized genes was performed using ShinyGO v0.85.1 (Ge et al., 2020). The 5,006 genes included in the Xenium Prime 5K dataset were used as background genes. GO enrichment was analyzed for Biological Process, Cellular Component, and Molecular Function categories. GO terms with an FDR of less than 0.05 were considered significantly enriched. The top enriched terms were visualized according to fold enrichment, number of genes, and −log10(FDR).

## Acknowledgements

This work was supported by JSPS KAKENHI (Grant Numbers 25K02372, 26H01604 to MH, 21K20853, 23K14203 and 26K18323 to TA); AMED (Grant Numbers 24wm0425005h0004 and 24ek0109764h0001 to MH), an Intramural Research Grant of NCNP (4–5, 4-6, 6-9 to MH); Japan Health Research Promotion Bureau (JH) under Research Fund (2024-D-01 to MH); Multilayered Stress Diseases (JPMXP1323015483 to MH) JST SPRING, (Grant Numbers JPMJSP2180 to KS); Tokumori Yasumoto Memorial Trust (to MH) and Takeda Science Foundation (Grants 2024049458 to TA). We thank T.O. and S.M. for fruitful discussions.

## Author contributions

Conceptualization: T.A., and M.H. Methodology: T.A., and M.H. Investigation: S.I., T.A., K.S. Formal analysis: S.I. Writing—original draft: S.I., T.A. and M.H. Writing—review and editing: All authors. Visualization: S.I., and T.A. Funding acquisition: T.A., and M.H. Supervision: T.A., M.S., and M.H.

## Data availability

All data supporting the findings of this study are provided within the paper and its Supplementary Information. Any additional information is available from the corresponding author upon reasonable request

## Code availability

No custom code was generated or used in this study.

## Figure legends

**Supplementary Figure 1.**
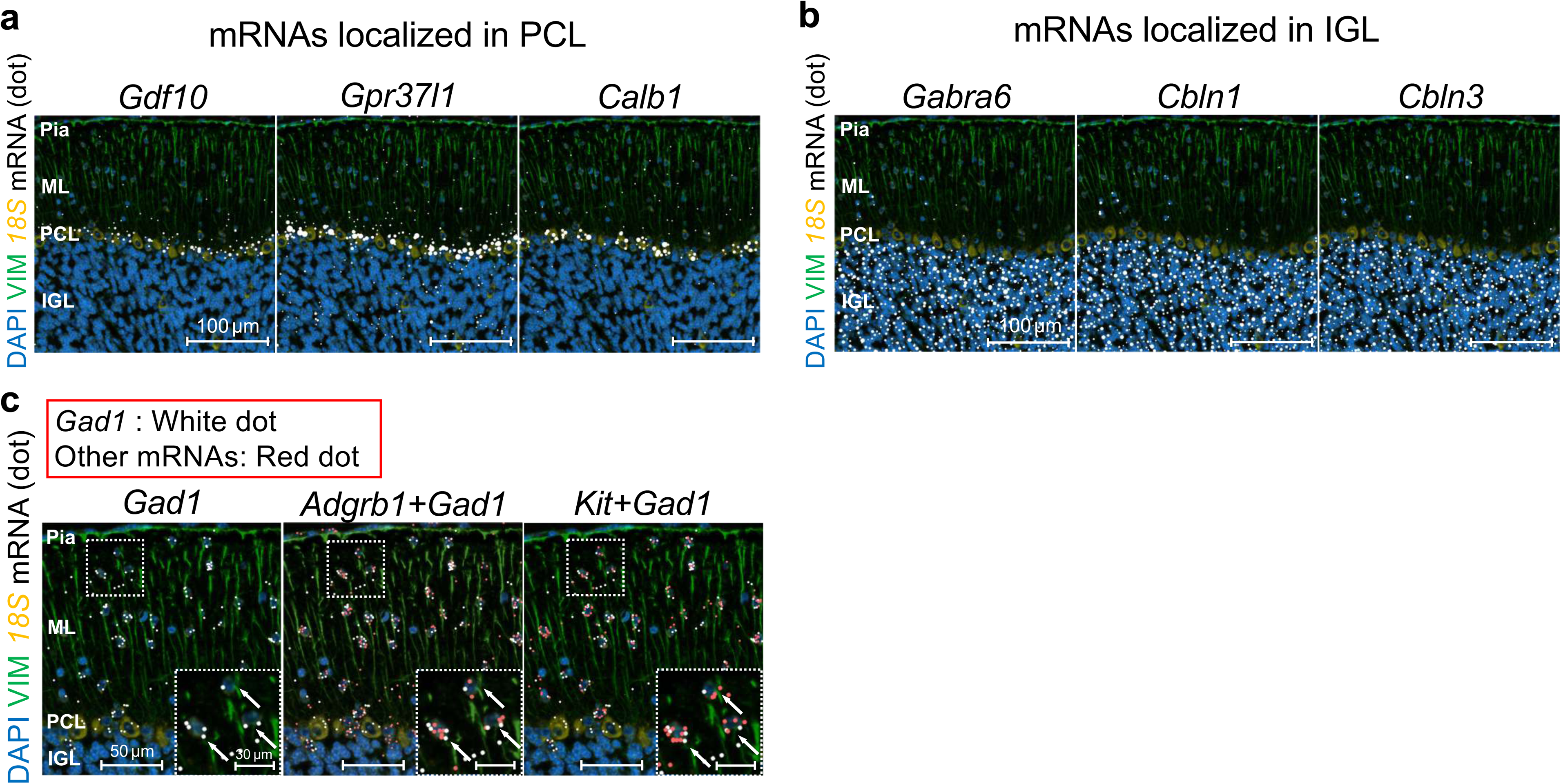
Layer- and cell body-associated mRNA distributions in the adult mouse cerebellar cortex detected by Xenium. (a) Representative Xenium images showing mRNAs predominantly localized in the Purkinje cell layer (PCL). *Gdf10*, *Gpr37l1*, and *Calb1* mRNA spots are shown in white. DAPI, VIMENTIN (VIM), and *18S* rRNA signals are shown in blue, green, and orange, respectively. Scale bars: 100 µm. (b) Representative Xenium images showing mRNAs predominantly localized in the inner granule cell layer (IGL). *Gabra6*, *Cbln1*, and *Cbln3* mRNA spots are shown in white. DAPI, VIM, and *18S* rRNA signals are shown in blue, green, and orange, respectively. Scale bars: 100 µm. (c) Representative Xenium images showing mRNAs associated with cell bodies in the molecular layer (ML). *Gad1* mRNA spots are shown in white, and other mRNA spots are shown in red. *Adgrb1* and *Kit* mRNA spots were visualized together with *Gad1* mRNA spots to identify genes detected in or near *Gad1*-positive ML cell bodies. Insets show magnified views of the boxed regions. Scale bars: 50 µm and 30 µm.

**Supplementary Figure 2.**
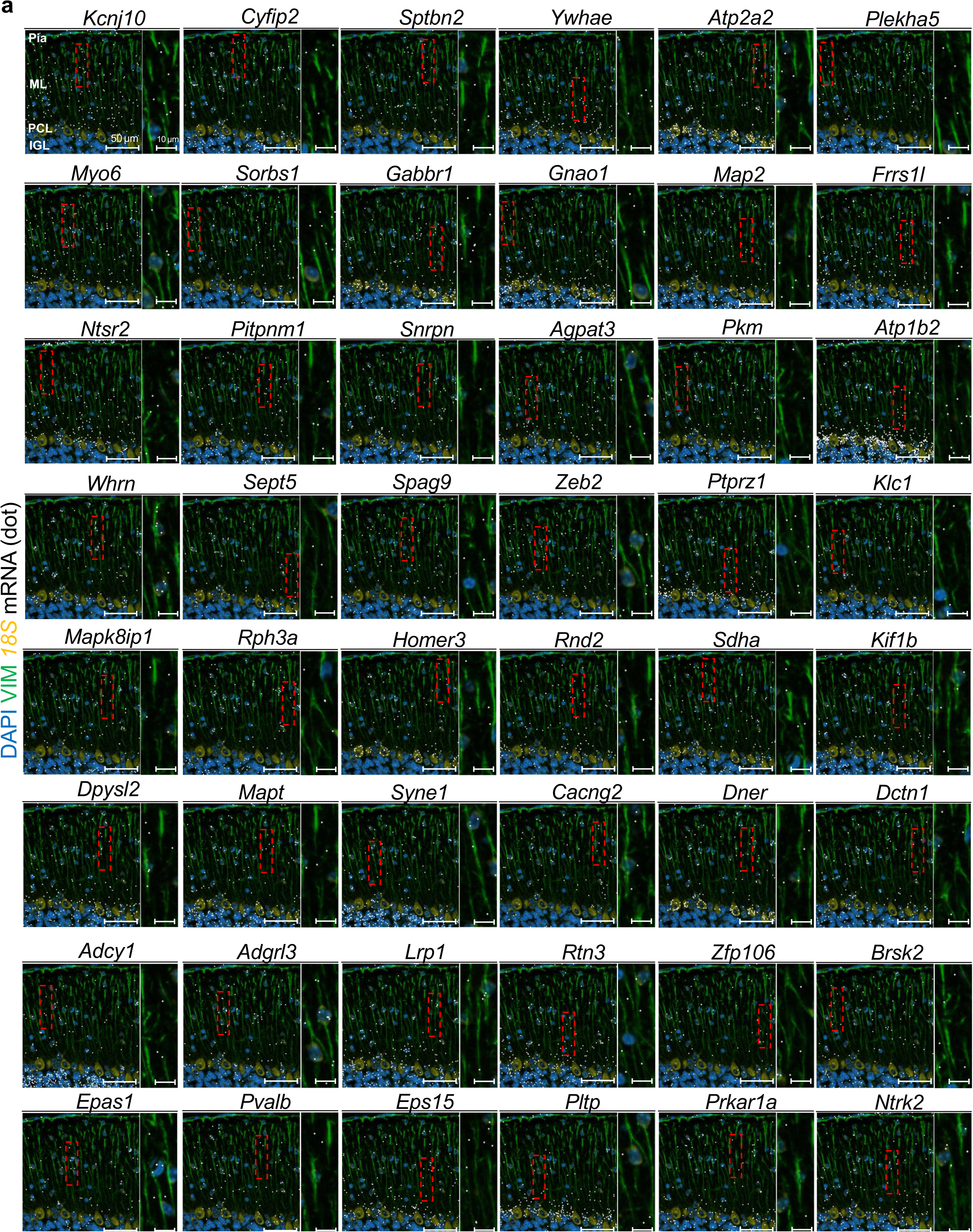
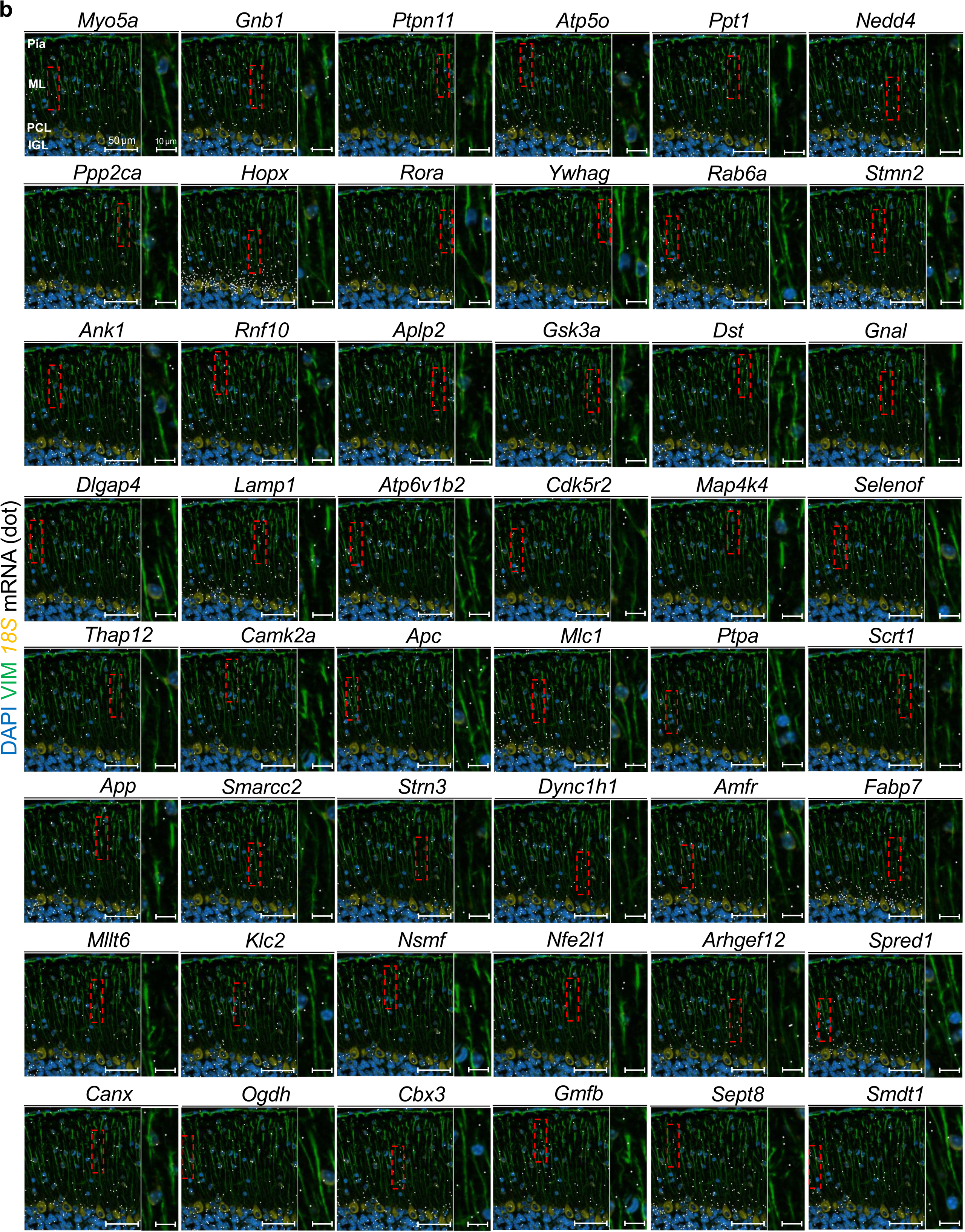
Additional candidate process-localized mRNAs detected in the adult cerebellar molecular layer. (a, b) Additional Xenium images showing candidate process-localized mRNAs detected in the adult mouse cerebellar molecular layer (ML). mRNA spots for each indicated gene are shown in white. DAPI, VIMENTIN (VIM), and 18S rRNA signals are shown in blue, green, and orange, respectively. Red dashed boxes indicate representative ML regions where mRNA spots were detected among densely distributed VIM-positive processes. Magnified images of the boxed regions are shown to the right of each panel. Scale bars: 50 µm and 10 µm.

**Supplementary Figure 3.**
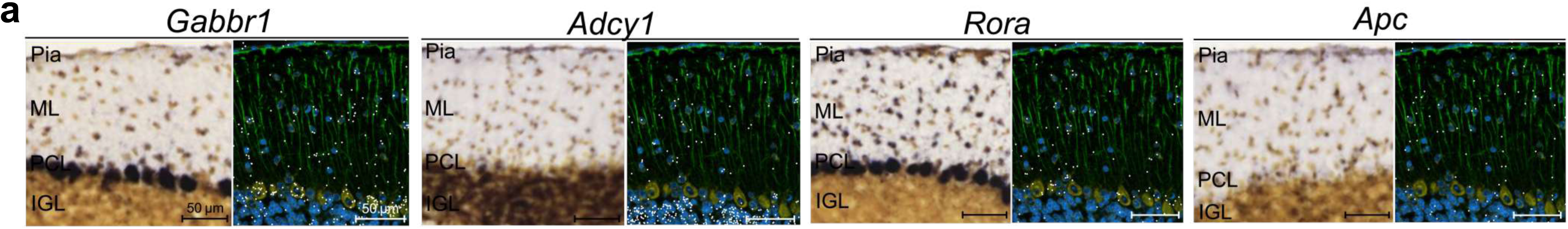
Candidate process-localized genes not clearly supported by Allen Brain Atlas ISH data. (a) Comparison of Allen Brain Atlas *in situ* hybridization (ISH) images and Xenium images for four candidate process-localized genes for which clear ML localization or process-like staining patterns were not observed in the available Allen Brain Atlas ISH images. For each gene, the Allen Brain Atlas ISH image is shown on the left, and the corresponding Xenium image is shown on the right. In the Xenium images, mRNA spots for each indicated gene are shown in white. DAPI, VIMENTIN (VIM), and *18S* rRNA signals are shown in blue, green, and orange, respectively. Scale bars: 50 µm.

**Supplementary Figure 4.**
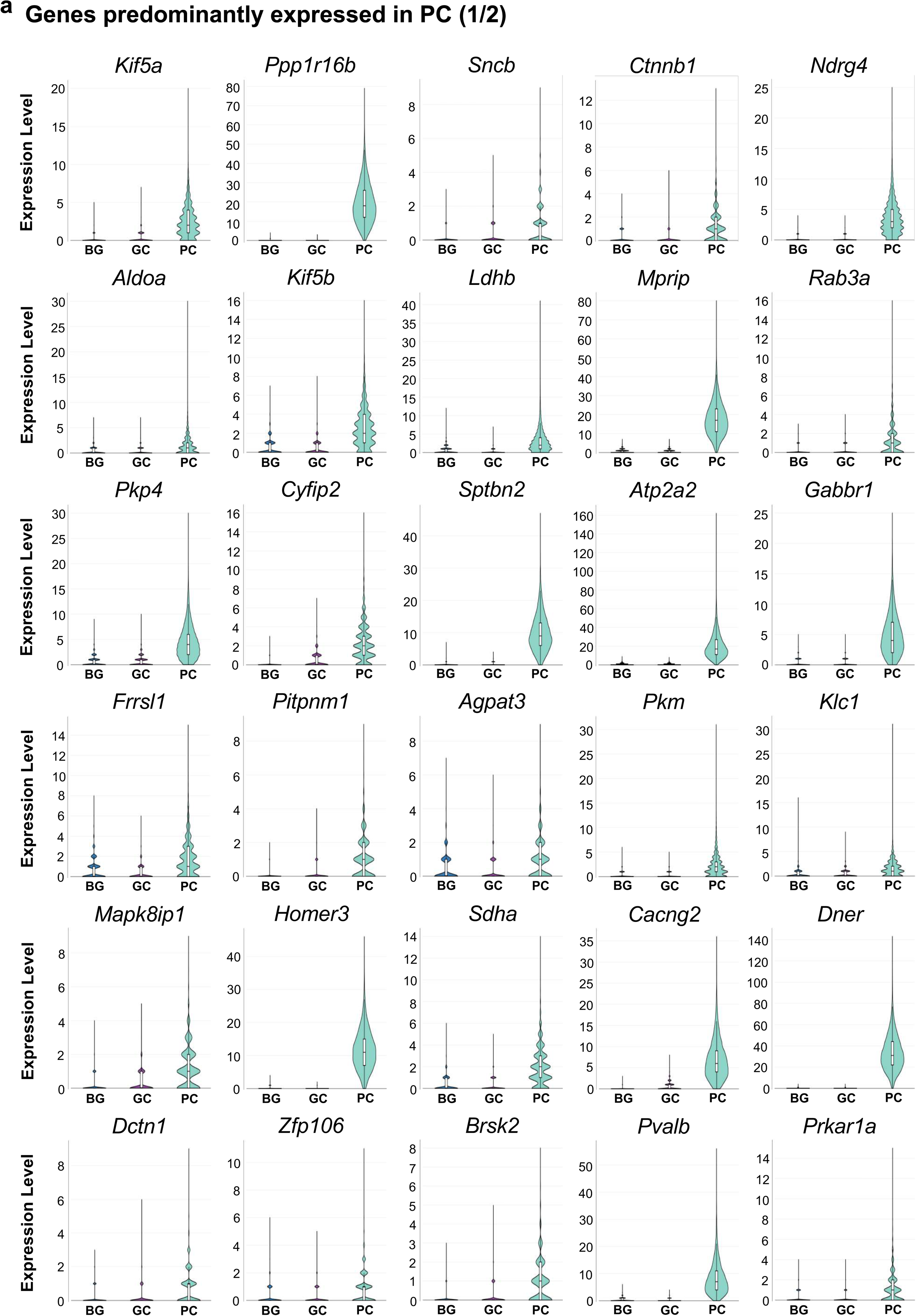
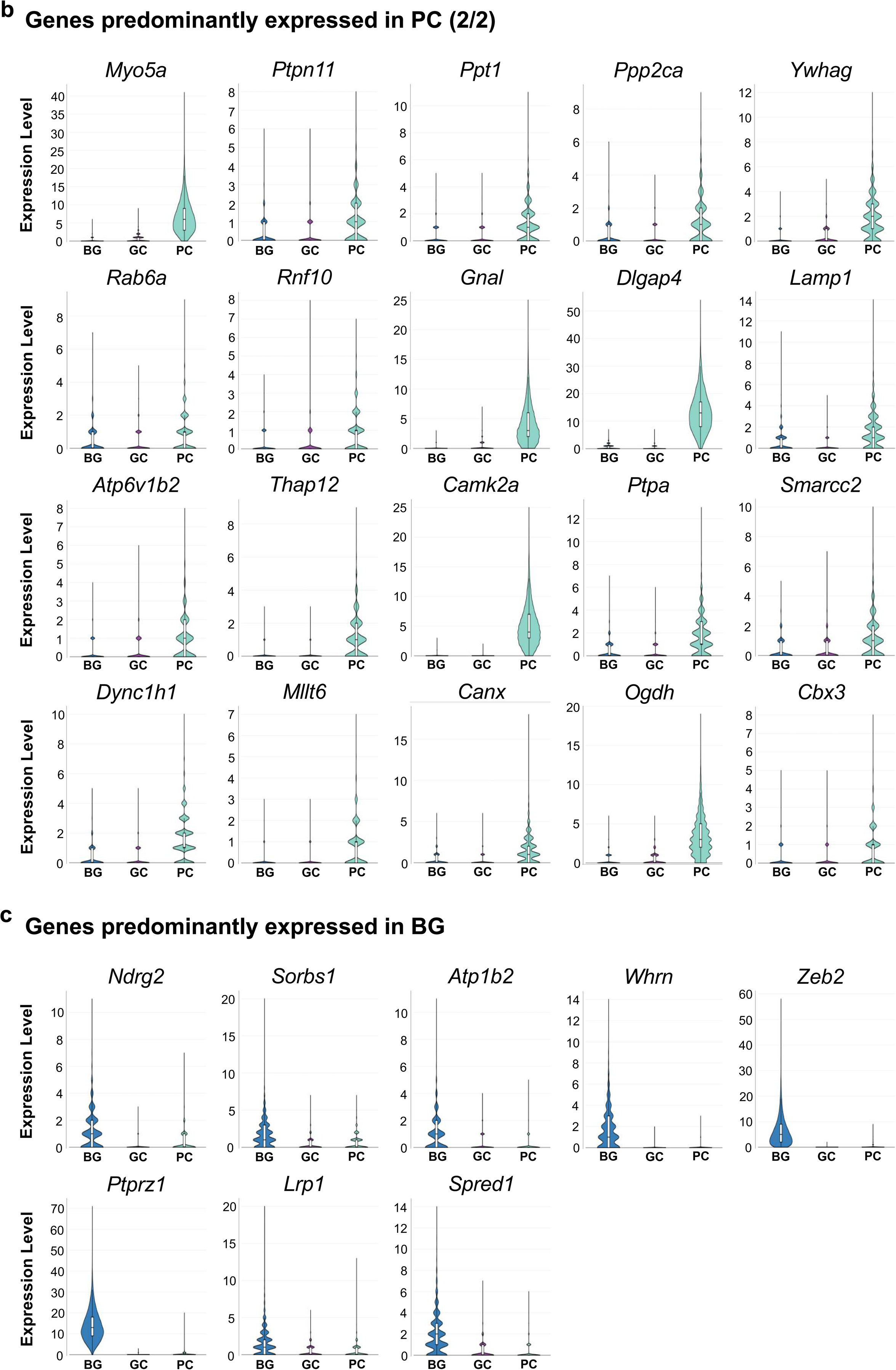
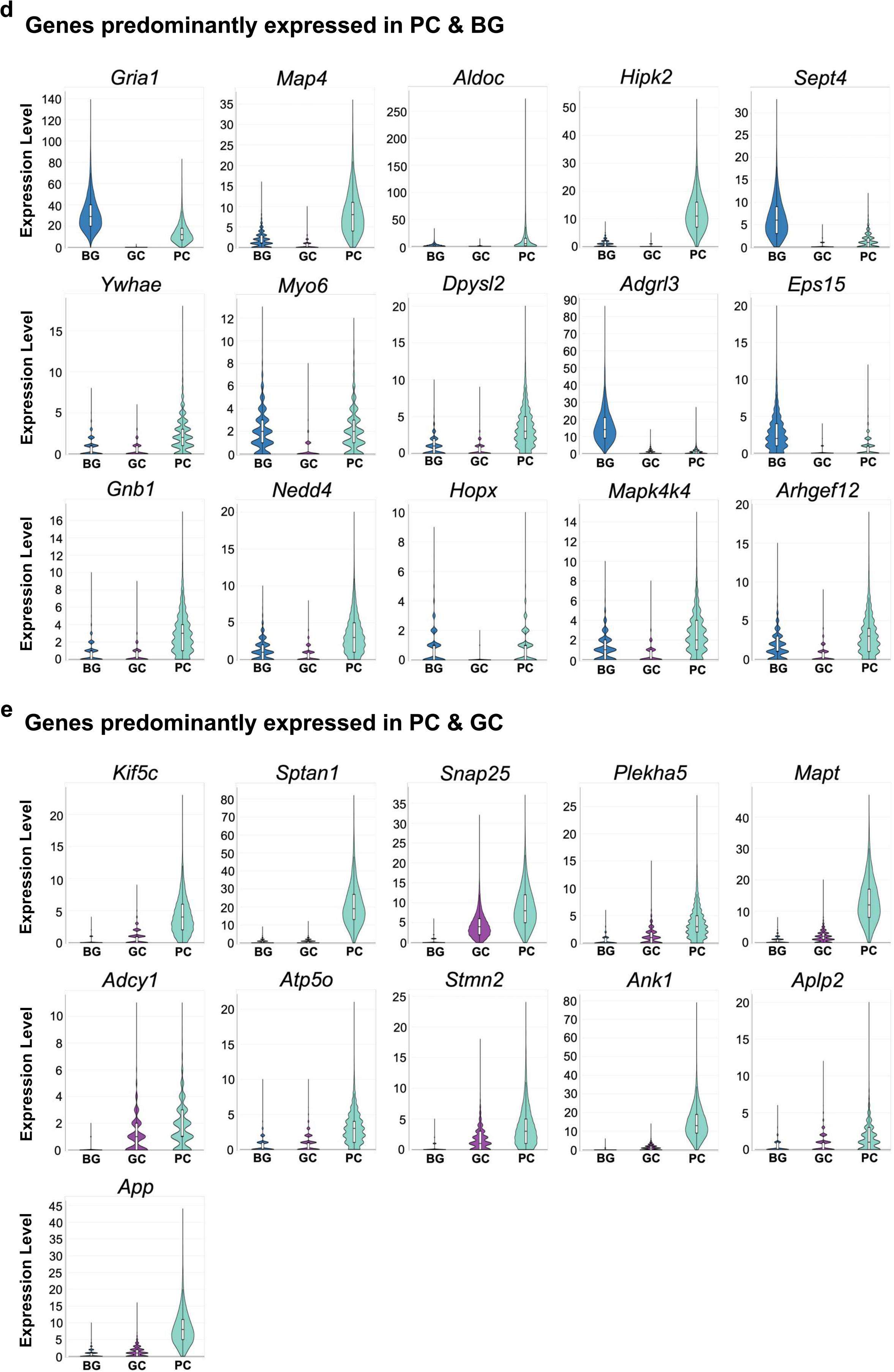
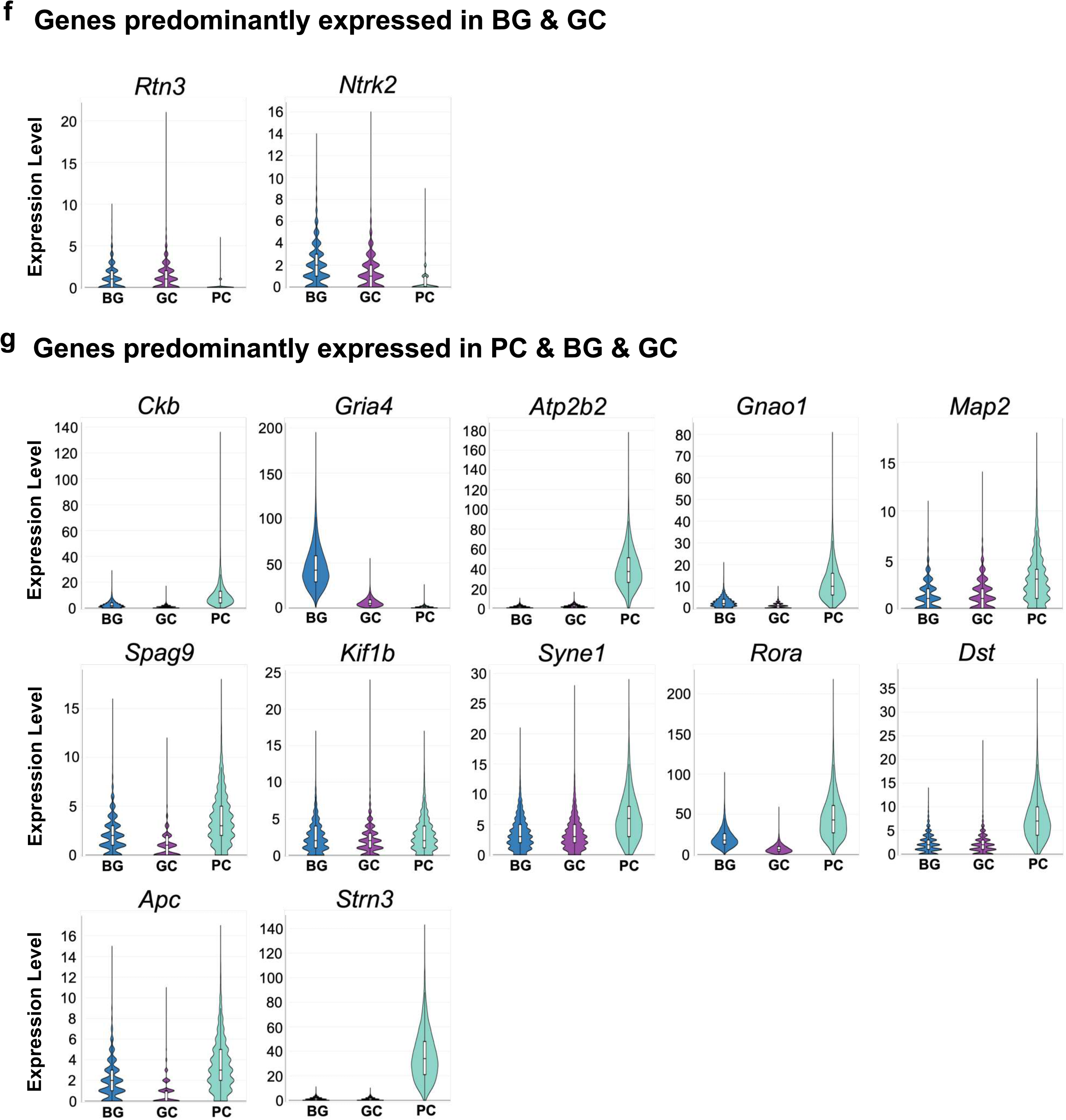
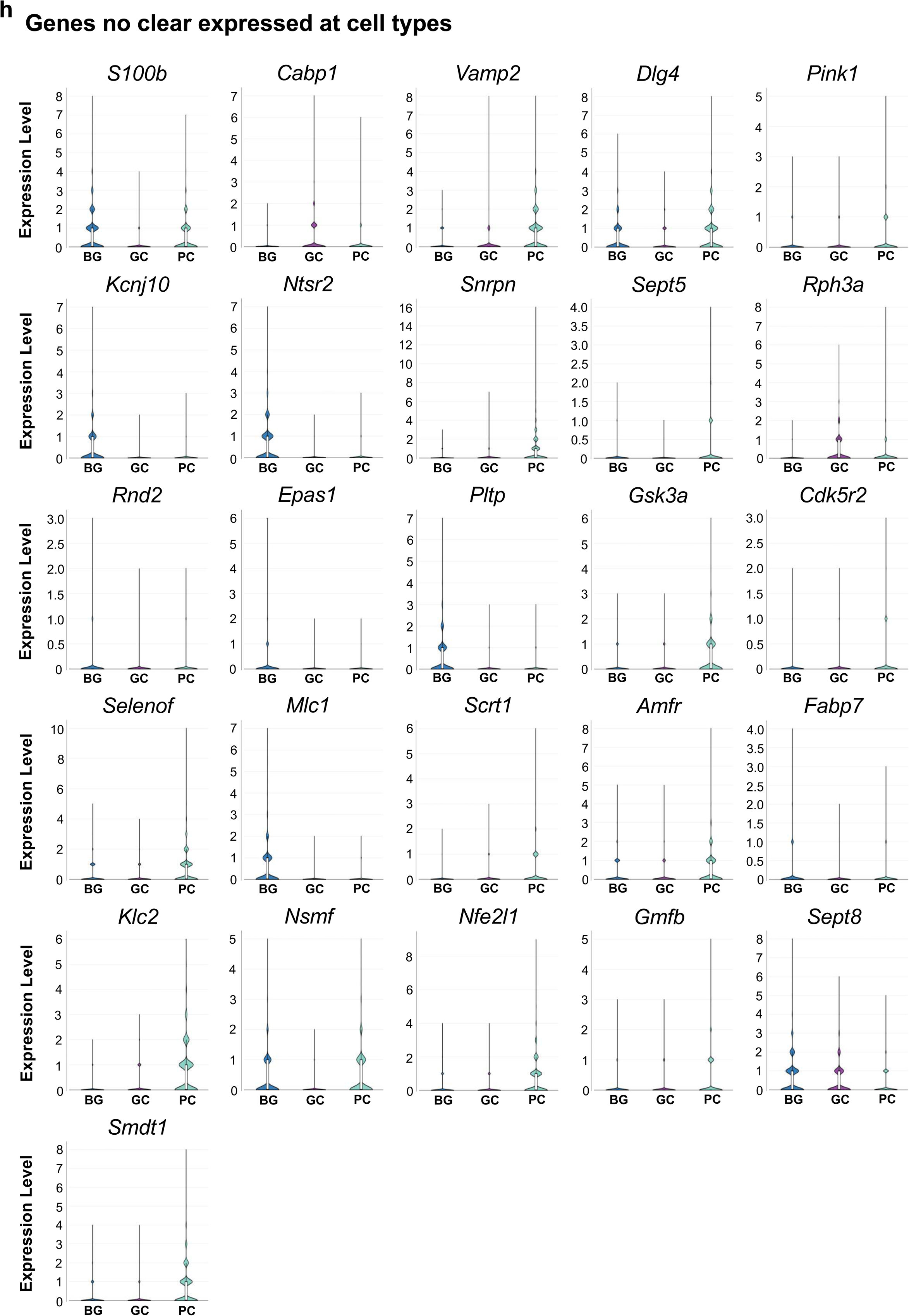
Cell-type expression profiles of candidate process-localized genes in a published cerebellar snRNA-seq atlas. (a–h) Violin plots showing the expression profiles of candidate process-localized genes in Bergmann glial cell (BG), granule cell (GC), and Purkinje cell (PC) clusters from the published adult mouse cerebellar single-nucleus RNA-seq (snRNA-seq) atlas reported by Kozareva et al. Genes were classified according to their cell-type expression patterns based on median unique molecular identifier (UMI) counts in each cluster. Genes with a median UMI count of 1.0 or higher in a given cell-type cluster were considered to be expressed in that cell type. Candidate genes were categorized as genes expressed in PC (a, b), BG (c), PC and BG (d), PC and GC (e), BG and GC (f), PC, BG, and GC (g), or genes with no clear expression in BG, GC, or PC clusters (h).

## References

Ango, F., Wu, C., Van der Want, J. J., Wu, P., Schachner, M., & Huang, Z. J. (2008). Bergmann glia and the recognition molecule CHL1 organize GABAergic axons and direct innervation of Purkinje cell dendrites. PLoS Biology, 6(4), e103. 10.1371/journal.pbio.0060103

Barton, S. K., Gregory, J. M., Chandran, S., & Turner, B. J. (2019). Could an impairment in local translation of mRNAs in glia be contributing to pathogenesis in ALS? Frontiers in Molecular Neuroscience, 12, 124. 10.3389/fnmol.2019.00124

Buxbaum, A. R., Haimovich, G., & Singer, R. H. (2015). In the right place at the right time: Visualizing and understanding mRNA localization. Nature Reviews Molecular Cell Biology, 16(2), 95–109. 10.1038/nrm3918

Cioni, J.-M., Koppers, M., & Holt, C. E. (2018). Molecular control of local translation in axon development and maintenance. Current Opinion in Neurobiology, 51, 86–94. 10.1016/j.conb.2018.02.025

Consalez, G. G., & Hawkes, R. (2013). The compartmental restriction of cerebellar interneurons. Frontiers in Neural Circuits, 6, 123. 10.3389/fncir.2012.00123

Dalla Costa, I., Buchanan, C. N., Zdradzinski, M. D., Sahoo, P. K., Smith, T. P., Thames, E., Kar, A. N., & Twiss, J. L. (2021). The functional organization of axonal mRNA transport and translation. Nature Reviews Neuroscience, 22(2), 77–91. 10.1038/s41583-020-00407-7

Dargan, R., Mikheenko, A., Johnson, N. L., Packer, B. E., Li, Z., Craig, E. J., Sarbanes, S. L., Bereda, C., Mehta, P. R., Keuss, M. J., Nalls, M. A., Qi, Y. A., Weller, C. A., Fratta, P., & Ryan, V. H. (2025). Altered mRNA transport and local translation in i³Neurons with RNA-binding protein knockdown. Nucleic Acids Research, 53(14), gkaf709. 10.1093/nar/gkaf709

De Zeeuw, C. I., & Hoogland, T. M. (2015). Reappraisal of Bergmann glial cells as modulators of cerebellar circuit function. Frontiers in Cellular Neuroscience, 9, 246. 10.3389/fncel.2015.00246

Doyle, M., & Kiebler, M. A. (2011). Mechanisms of dendritic mRNA transport and its role in synaptic tagging. The EMBO Journal, 30(17), 3540–3552. 10.1038/emboj.2011.278

Ge, S. X., Jung, D., & Yao, R. (2020). ShinyGO: A graphical gene-set enrichment tool for animals and plants. Bioinformatics, 36(8), 2628–2629. 10.1093/bioinformatics/btz931

Gumy, L. F., Yeo, G. S. H., Tung, Y.-C. L., Zivraj, K. H., Willis, D., Coppola, G., Lam, B. Y. H., Twiss, J. L., Holt, C. E., & Fawcett, J. W. (2011). Transcriptome analysis of embryonic and adult sensory axons reveals changes in mRNA repertoire localization. RNA, 17(1), 85–98. 10.1261/rna.2386111

Holt, C. E., & Schuman, E. M. (2013). The central dogma decentralized: New perspectives on RNA function and local translation in neurons. Neuron, 80(3), 648–657. 10.1016/j.neuron.2013.10.036

Janesick, A., Shelansky, R., Gottscho, A. D., Wagner, F., Williams, S. R., Rouault, M., Beliakoff, G., Morrison, C. A., Oliveira, M. F., Sicherman, J. T., Kohlway, A., Abousoud, J., Drennon, T. Y., Mohabbat, S. H., 10x Development Teams, & Taylor, S. E. B. (2023). High resolution mapping of the tumor microenvironment using integrated single-cell, spatial and in situ analysis. Nature Communications, 14, 8353. 10.1038/s41467-023-43458-x

Jung, H., & Holt, C. E. (2011). Local translation of mRNAs in neural development. Wiley Interdisciplinary Reviews: RNA, 2(1), 153–165. 10.1002/wrna.53

Kozareva, V., Martin, C., Osorno, T., Rudolph, S., Guo, C., Vanderburg, C., Nadaf, N., Regev, A., Regehr, W. G., & Macosko, E. Z. (2021). A transcriptomic atlas of mouse cerebellar cortex comprehensively defines cell types. Nature, 598(7879), 214–219. 10.1038/s41586-021-03220-z

Lee, B. H., Bae, S.-W., Shim, J. J., Park, S. Y., & Park, H. Y. (2016). Imaging single-mRNA localization and translation in live neurons. Molecules and Cells, 39(12), 841–846. 10.14348/molcells.2016.0277

Lein, E. S., Hawrylycz, M. J., Ao, N., Ayres, M., Bensinger, A., Bernard, A., Boe, A. F., Boguski, M. S., Brockway, K. S., Byrnes, E. J., Chen, L., Chen, L., Chen, T.-M., Chin, M. C., Chong, J., Crook, B. E., Czaplinska, A., Dang, C. N., Datta, S., … Jones, A. R. (2007). Genome-wide atlas of gene expression in the adult mouse brain. Nature, 445(7124), 168–176. 10.1038/nature05453

Leto, K., Arancillo, M., Becker, E. B. E., Buffo, A., Chiang, C., Ding, B., Dobyns, W. B., Dusart, I., Haldipur, P., Hatten, M. E., Hoshino, M., Joyner, A. L., Kano, M., Kilpatrick, D. L., Koibuchi, N., Marino, S., Martinez, S., Millen, K. J., Millner, T. O., … Hawkes, R. (2016). Consensus paper: Cerebellar development. The Cerebellum, 15(6), 789–828. 10.1007/s12311-015-0724-2

Lin, A. C., & Holt, C. E. (2007). Local translation and directional steering in axons. The EMBO Journal, 26(16), 3729–3736. 10.1038/sj.emboj.7601808

Meservey, L. M., Topkar, V. V., & Fu, M.-M. (2021). mRNA transport and local translation in glia. Trends in Cell Biology, 31(6), 419–423. 10.1016/j.tcb.2021.03.006

Middleton, S. A., Eberwine, J., & Kim, J. (2018). Comprehensive catalog of dendritically localized mRNA isoforms from sub-cellular sequencing of single mouse neurons [Preprint]. bioRxiv. 10.1101/278648

Mofatteh, M. (2020). mRNA localization and local translation in neurons. AIMS Neuroscience, 7(3), 299–310. 10.3934/Neuroscience.2020016

Ohashi, R., & Shiina, N. (2020). Cataloguing and selection of mRNAs localized to dendrites in neurons and regulated by RNA-binding proteins in RNA granules. Biomolecules, 10(2), 167. 10.3390/biom10020167

Sakers, K., Lake, A. M., Khazanchi, R., Ouwenga, R., Vasek, M. J., Dani, A., & Dougherty, J. D. (2017). Astrocytes locally translate transcripts in their peripheral processes. Proceedings of the National Academy of Sciences of the United States of America, 114(19), E3830–E3838. 10.1073/pnas.1617782114

Singer, A., Trigo, F., Vinel, L., Gruere, O., Llano, I., & Oheim, M. (2025). A first morphological and electrophysiological characterization of Fañanas cells of the mouse cerebellum. The Journal of Physiology, 603(4), 855–871. 10.1113/JP285949

Sotelo, C. (2015). Molecular layer interneurons of the cerebellum: Developmental and morphological aspects. The Cerebellum, 14(5), 534–556. 10.1007/s12311-015-0648-x

Sunkin, S. M., Ng, L., Lau, C., Dolbeare, T., Gilbert, T. L., Thompson, C. L., Hawrylycz, M., & Dang, C. (2013). Allen Brain Atlas: An integrated spatio-temporal portal for exploring the central nervous system. Nucleic Acids Research, 41(D1), D996–D1008. 10.1093/nar/gks1042

Suyama, K., Adachi, T., Mizuno, M., Ji, K., Isogai, E., Hasegawa, I., Nishitani, K., Sone, M., Miyashita, S., Owa, T., & Hoshino, M. (in press). Molecular characteristics and differentiation control mechanisms of Bergmann glia-like progenitors in the postnatal mouse cerebellum. Glia. 10.1002/glia.70160

Taylor, A. M., Berchtold, N. C., Perreau, V. M., Tu, C. H., Jeon, N. L., & Cotman, C. W. (2009). Axonal mRNA in uninjured and regenerating cortical mammalian axons. The Journal of Neuroscience, 29(15), 4697–4707. 10.1523/JNEUROSCI.6130-08.2009

Turner-Bridger, B., Caterino, C., & Cioni, J.-M. (2020). Molecular mechanisms behind mRNA localization in axons. Open Biology, 10(9), 200177. 10.1098/rsob.200177

Williams, C. G., Lee, H. J., Asatsuma, T., Vento-Tormo, R., & Haque, A. (2022). An introduction to spatial transcriptomics for biomedical research. Genome Medicine, 14, 68. 10.1186/s13073-022-01075-1

Xing, L., & Bassell, G. J. (2013). mRNA localization: An orchestration of assembly, traffic and synthesis. Traffic, 14(1), 2–14. 10.1111/tra.12004

Zappulo, A., van den Bruck, D., Ciolli Mattioli, C., Franke, V., Imami, K., McShane, E., Moreno-Estelles, M., Calviello, L., Filipchyk, A., Peguero-Sanchez, E., Müller, T., Woehler, A., Birchmeier, C., Merino, E., Rajewsky, N., Ohler, U., Mazzoni, E. O., Selbach, M., Akalin, A., & Chekulaeva, M. (2017). RNA localization is a key determinant of neurite-enriched proteome. Nature Communications, 8, 583. 10.1038/s41467-017-00690-6

